# Using gnotobiotic mice to decipher effects of gut microbiome repair in undernourished children on tuft and goblet cell function

**DOI:** 10.1101/2025.10.02.680046

**Authors:** Yi Wang, Hao-Wei Chang, Jiye Cheng, Daniel M. Webber, Hannah M. Lynn, Matthew C. Hibberd, Clara Kao, Ishita Mostafa, Tahmeed Ahmed, Michael J. Barratt, Jeffrey I. Gordon

**Affiliations:** Edison Family Center for Genome Sciences and Systems Biology, Washington University School of Medicine, St. Louis, MO 63110 USA; Newman Center for Gut Microbiome and Nutrition Research, Washington University School of Medicine, St. Louis, MO 63110 USA; Department of Pathology and Immunology, Washington University School of Medicine, St. Louis, MO 63110 USA; International Centre for Diarrhoeal Disease Research, Bangladesh (icddr,b), Dhaka 1212, Bangladesh

**Keywords:** childhood undernutrition, gut microbiome-directed therapeutic foods, barrier function, tuft cells, goblet cells, gnotobiotic mice

## Abstract

Studies have implicated perturbations in the postnatal development of the gut microbiome as a contributing factor to childhood undernutrition. Compared to a standard ready-to-use supplementary food, a microbiome-directed complementary food (MDCF-2) designed to repair these perturbations produced superior improvements in ponderal and linear growth in clinical trials of Bangladeshi children with moderate acute malnutrition. Here, ‘reverse translation’ experiments are performed where intact fecal microbiomes collected from trial participants prior to and at the end of treatment are introduced into female gnotobiotic mice just after delivery of their pups. Pups received diets designed to resemble those consumed by children in the trials to recreate “unrepaired” and “repaired” gut ecosystems. Analyses of the abundances of bacterial strains (metagenome-assembled genomes), their expressed genes and metabolic products, combined with assessments of ponderal growth and intestinal epithelial lineage transcriptomes (single-nucleus RNA-Seq with follow-up immunocytochemistry) disclosed effects of MDCF-2 associated microbiome repair that cannot be determined, in part because ‘no treatment’ control arms cannot be ethically incorporated into these trials. Specifically, microbiome repair in these mice produced significant increases in ponderal growth, changes microbial gene expression consistent with a less virulent gut ecosystem and changes in expression of (i) components of gut epithelial cell junctions in the enterocytic and goblet cell lineages, (ii) pathways for synthesis and secretion of eicosanoid immune effectors in chemosensory tuft cells, and (iii) goblet cell pathways involved in glycosylation and secretion of mucin. Experiments of the type described can help formulate and test hypotheses about how microbiome repair affects host biology.

**SIGNIFICANCE STATEMENT:** Undernutrition is a global health problem. Recent clinical trials of a gut microbiome-directed complementary food (MDCF-2) designed to repair the perturbed gut microbiomes of undernourished Bangladesh children produced superior growth outcomes versus a standard nutritional supplement. Given ethical considerations and tissue sampling constraints associated with these types of studies, we colonized gnotobiotic mice postnatally with microbiome samples obtained from trial participants before and after treatment to model “unrepaired” and “repaired” gut ecosystems. Using a multi-omics approach, we uncover heretofore unappreciated changes in expressed chemosensory tuft cell, mucus-producing goblet cell and absorptive enterocytic functions and interactions accompanying microbiome repair. Extending microbiome clinical trials back to preclinical models (‘reverse translation’) provides mechanistic insights that can inform design/interpretation of future interventions.

## INTRODUCTION

Childhood undernutrition represents a pressing global health challenge. Epidemiologic studies indicate that factors in addition to food insecurity contribute to pathogenesis. One such factor is a disruption in the postnatal assembly and functional maturation of the gut microbiome in undernourished infants/children. This process is largely completed during the first 2-3 years after birth in healthy individuals (1). Emerging evidence suggests that proper coordinated co-development of the gut microbiome and host organ systems is an important contributor to healthy postnatal growth (2–4).

A randomized controlled clinical trial of 12-18-month-old Bangladeshi children with primary moderate acute malnutrition (MAM) has demonstrated that dietary supplementation with a microbiome-directed complementary food (MDCF-2) designed to repair their gut communities significantly improved their ponderal as well as linear growth compared to a commonly used ready-to-use supplementary food (RUSF) formulation, even though the RUSF had 15% higher caloric density than the MDCF (5). DNA and mRNA analyses of fecal samples, serially collected during the course 3-month long-period of MDCF-2 intervention, disclosed that two *Prevotella copri* strains, whose abundances were positively associated with improvements in participants’ ponderal growth, were the predominant source of transcripts encoding enzymes involved in the metabolism of MDCF polysaccharides (6). A similar result was obtained in a separate randomized controlled clinical trial of 12-18-month-old Bangladeshi children who had presented with severe acute malnutrition (SAM), had undergone a hospital-based initial nutritional resuscitation protocol that was not designed to repair their microbiome and were left with MAM (i.e., they had post-SAM MAM). Compared to RUSF, MCDF-2 treatment in both trials was accompanied by significantly greater changes in the levels of multiple plasma protein biomarkers and mediators of musculoskeletal and CNS development, immune function and metabolic regulation (7).

Our previous work utilized a gnotobiotic mouse model that involved dam-to-pup transmission of a defined community of age- and growth-associated bacterial strains cultured from Bangladeshi infants/children to dissect the mechanisms underlying MDCF-2’s effects (8). We found that inclusion of *P. copri* in the bacterial consortium significantly increased postnatal weight gain in a MDCF-2-dependent fashion and significantly increased the metabolism of polysaccharides deemed to be key bioactive components of MDCF-2 (8). Moreover, single nucleus RNA-sequencing (snRNA-Seq), combined with targeted mass spectrometric analysis of intestinal segments disclosed that the presence of *P. copri* had a marked effect on metabolism in the predominant gut epithelial cell lineage (enterocytes) (8).

These mechanistic studies used defined culture collections that might not fully capture the complexity of the intact gut luminal ecosystem. In the current report, we compare offspring of gnotobiotic dams colonized with intact uncultured microbiota from children in the upper quartile of clinical response to MDCF-2 in the clinical trial of children with primary MAM. Our goal was to compare the outcomes of MDCF-2 treatment using multi-omic analyses to those of a hypothetical scenario in which the undernourished participants had not received any treatment.

## RESULTS

### Experimental Design

**Fig. 1A** summarizes the experimental design. Intact uncultured fecal samples that had been stored at -80 °C since their collection were selected from two participants in the randomized controlled clinical study of Bangladeshi children with primary MAM. Among trial participants, these children fell within the upper quartile of ponderal growth response following MDCF-2 treatment, as measured by changes in weight-for-length z-score (WLZ). They also exhibited increases in the abundances of bacterial taxa that were positively associated with WLZ and decreases in taxa negatively associated with WLZ (5). Two fecal samples were used from each child: one collected just before initiating MDCF-2 treatment and one obtained at the conclusion of the 3-month intervention with this therapeutic food. These samples were designated “unrepaired” (pre-treatment) and “repaired” (post-treatment) microbial communities. Three custom diets were formulated for these experiments (**Dataset S1**): (i) a ‘weaning diet’ reflective of that consumed by 12-18-month-old Bangladeshi children living in the Mirpur urban slum where the randomized controlled clinical trials had been conducted, (ii) a diet representative of the diet consumed by weaned 18-month-old children living in Mirpur (Mirpur-18 diet), and (iii) MCDF-2. Diets were irradiated to ensure their sterility.

**Fig. 1.**
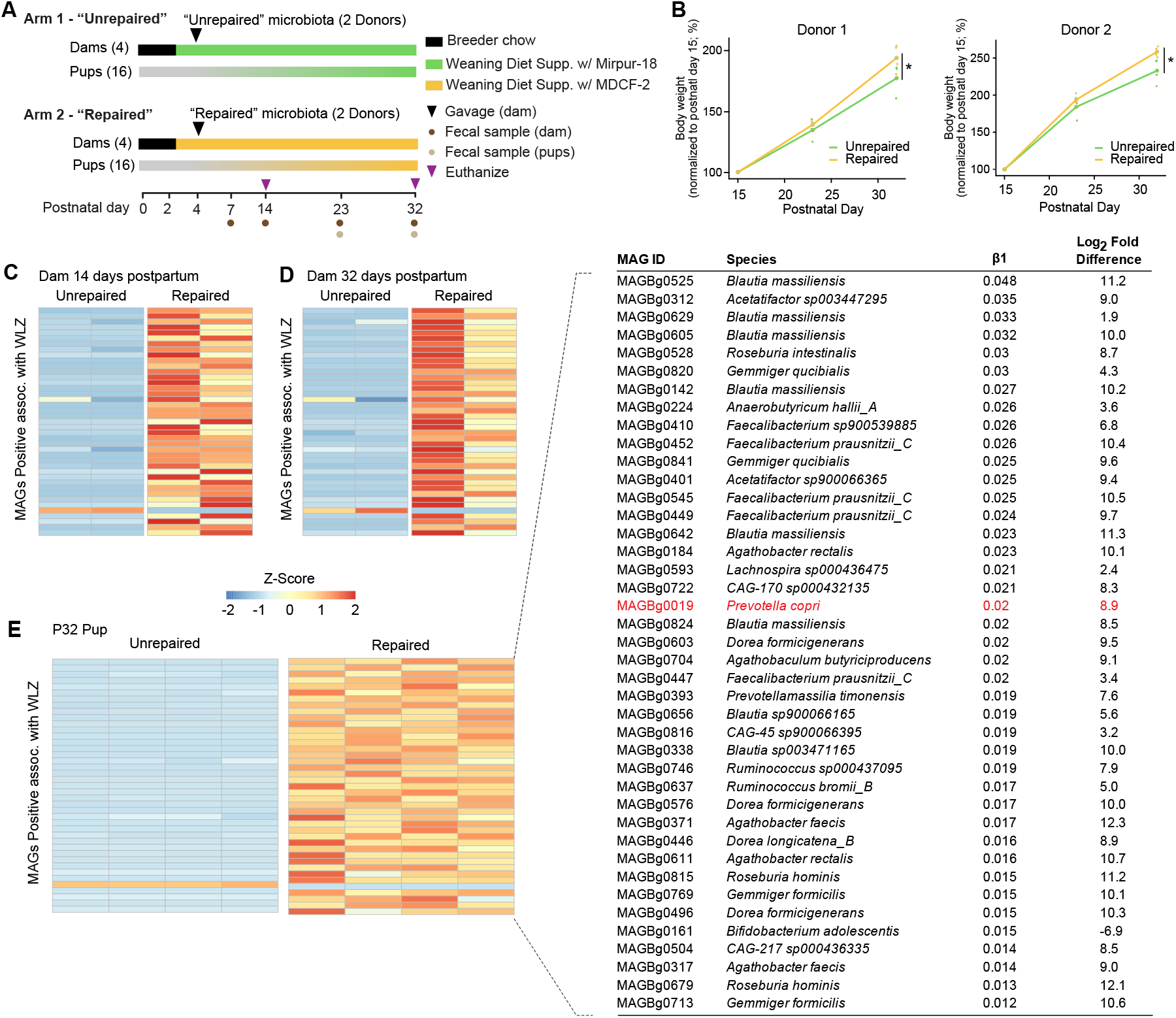
MAG and microbial RNA-seq analyses. **A**. Experimental Design. **B**. Line plots illustrating relative weight changes from postnatal day 15 (P15) to P32, using P15 as the reference point for comparison. Data are presented for both donor 1 and donor 2 experiments. Error bars represent standard deviation. **\*** indicates statistical significance with *P* < 0.05. **C-E**. Heatmaps representing the relative abundance of colonized MAGs positively associated with WLZ in postpartum day 14 (C), postpartum day 32 dams (D), and postnatal day 32 pups (E) for the donor 1 experiment. Z-score was calculated by centering and scaling the log-transformed relative abundances of each MAG across all samples. Each column represents data from an individual animal. The inset in **E** provides detailed information about the WLZ-associated MAGs shown in the heatmap, including MAG ID, assigned species, β1 coefficient from the linear mixed effects model [WLZ ∼ β1(MAG) + β2(study week) + 1|PID] indicating the strength of WLZ association, the Log2-fold difference in MAG abundance between “unrepaired” and “repaired” arms. The MAGs are ordered by descending β1 coefficient. *P. copri* is highlighted in red.

Eight germ-free female C57BL/6J mice that had been maintained on standard breeder chow during their pregnancy delivered a total of 36 pups; nine pups were randomly assigned on postpartum day 1 (P1) to each of four groups of dually housed dams that had been switched to the weaning diet. On postpartum day 2 (P2) this weaning diet was supplemented for 2 groups of dams with Mirpur-18 while the other 2 groups received the weaning diet supplemented with MDCF-2 (**Dataset S1**). Two days later (postpartum day 4), each co-housed dam consuming the Mirpur-18-supplmented weaning diet was gavaged with a pre-treatment ‘unrepaired’ fecal microbiome sample from one or the other of the two donor children with MAM, while a separate of co-housed dams consuming the weaning diet supplemented with MDCF-2 received the corresponding post-treatment ‘repaired’ community from the same child (total of 4 experimental groups). Thus, maternal-to-pup transfer of these donor communities occurred while the pups experienced a diet sequence that first began with exclusive milk feeding (from the nursing dam) followed by a weaning period where pups consumed the dam’s milk plus the ‘weaning diet’ supplemented with Mirpur-18 or with MDCF-2 (**Dataset S1**). Five pups were euthanized on postnatal day 14 (P14) from each group. The remaining offspring were maintained with the dams until they were euthanized on P32 (**Fig. 1A**). To control for diet effects, we included two additional germ-free control groups that received either the Mirpur-18–supplemented or the MDCF-2–supplemented weaning diet (n=2 dams with 11 pups per group) .

### Effects on ponderal growth

Mice that received the ‘repaired’ microbiota from either donor demonstrated significantly greater increases in body weight from P15 to P32 than mice that had received ‘unrepaired’ microbiota from the same donor [*P*< 0.05; linear mixed-effects model (mixed effects: group and postnatal day; random effect: mouse); **Fig. 1B**]. As the weaning diet supplemented with MDCF-2 has 10% more energy than the Mipur-18-supplemented diet (**Dataset S1**), both dietary differences and microbiome configuration could have contributed to the weight gain phenotype. To disentangle these effects, we first assessed the effects of weaning diets supplemented with MDCF-2 versus Mirpur-18 on the ponderal growth of uncolonized (germ-free) mice. Germ-free pups fed the weaning diet supplemented with MDCF-2 exhibited a statistically significant increase in weight gain compared to those receiving the weaning diet supplemented with Mirpur-18 [*P*=0.014; linear mixed-effects model: relative weight gain ∼ diet × days of experiment + (1 | mouse)]. We subsequently pooled data from the donor 1- and donor 2-colonized and germ-free mice and applied a linear mixed-effects model [Relative_weight_gain ∼ Diet * Day_of_experiment + Microbiota * Day_of_experiment + (1|Mouse_ID)]. This analysis revealed that both the interaction term of microbiota and time of the experiment (*P*=5.34×10^-15^) and the interaction term of diet and time of the experiment (*P*=0.038) contributed significantly to the observed difference in weight gain. Together, these results indicated that both the MDCF-2-supplemented weaning diet and the repaired microbiome contribute to the observed growth enhancement.

### WLZ-associated metagenome-assembled genomes (MAGs) in repaired versus unrepaired microbiomes

We performed shotgun sequencing of DNAs isolated from (i) fecal samples serially collected from dams at postpartum days 7, 14, 23, and 32, (ii) fecal samples collected from their pups at P23 and P32 and (iii) cecal contents harvested from pups at the time of their euthanasia at P32 (**Dataset S2**). The resulting dataset was used to quantify the representation of 222 MAGs that we had previously found to be significantly positively (n=75) or negatively (n=147) associated with ponderal growth (WLZ) in children with primary MAM who participated in the clinical trial (6). Postpartum day 32 dams that received unrepaired or repaired fecal microbiomes from donor 1 contained 107 and 141 WLZ-associated MAGs, respectively (colonization criteria: TPM > 20 and prevalence > 40%). 97.7±5.4% (mean±SD) and 93.8±0.9% of these MAGs were also present in the cecal microbiomes of their P32 offspring (n=4 mice/treatment group; **Dataset S2C**). Postpartum day 32 dams colonized with donor 2’s unrepaired and repaired fecal communities contained 111 and 74 WLZ-associated MAGs, respectively, of which 94.4±1.5% and 86.1±7.2% were transmitted to their P32 offspring (n=4 mice/treatment group; **Dataset S2C**). 63.1% and 92.2% of the transmitted MAGs in donor 1 and donor 2 P32 pup cecal microbiomes were shared across the two experiments (highlighted in **Dataset S2D**). Volcano plots of the WLZ-association coefficient β1 and q-values (**SI Appendix, Fig. S1A**) demonstrate that bacterial strains (MAGs) most strongly associated with ponderal growth in the clinical trial had successfully colonized the dams.

This gnotobiotic dam-to-pup transmission model recapitulated the enhanced representation of positive WLZ-associated MAGs in repaired microbiomes; this was evident in dams at postpartum days 14 and 32, and in all of their pups at P32 in both the donor 1 and donor 2 experiments (**Fig. 1C-E, Dataset S2C**). In the clinical trial, *P. copri* MAG Bg0019 was a major source of MDCF-2-induced transcripts for carbohydrate-active enzymes (CAZYmes) involved in the metabolism of its bioactive glycan components (6). The abundance of this MAG, which is positively associated with WLZ (6), was significantly greater in the repaired compared to unrepaired microbiomes of dam-pup dyads that had received either the donor 1 or the donor 2 community (**Fig. 1E**).

### Transcriptional responses of small intestinal epithelial cell lineages to microbiome repair

To assess the intestinal epithelial response to microbiome repair along the crypt-to-villus and duodenal-to-ileal axes of the intestine, single-nucleus RNA sequencing (snRNA-Seq) was performed on duodenal, jejunal and ileal segments harvested from 3 animals in each of the four treatment groups (**Dataset S3**). Enterocytes, enteroendocrine cells, goblet cells, tuft cells, Paneth cells, transient amplifying and stem cells, plus intraepithelial lymphocytes were identified based on expression of known marker genes (**SI Appendix Fig. S1B**,**C**). Enterocytes were further classified into three subclusters corresponding to their location at the base, middle and tips of villi **(SI Appendix Fig. S1C**). Pseudobulk analysis (see *Methods*) was used to identify genes that were significantly differentially expressed in mice receiving unrepaired compared to repaired fecal microbiomes from donors 1 and 2 [threshold cutoffs: FDR-adjusted P-value (q-value) < 0.1; fold-difference in transcript levels > 1.5] (**Dataset S3B**). We then used the *Compass* algorithm (9) to assess the effects of MDCF-2-mediated microbiome repair on the differential expression of metabolic pathways in seven epithelial cell clusters identified in our snRNA-Seq datasets. Compass employs *in silico* flux balance analysis to predict metabolic reaction rates based on mass balance constraints and enzymatic capacity for each metabolic reaction curated in the Recon 2 reconstructed model of human metabolism (10). We focused on reactions inferred to have significantly altered flux with consistent directionality of change across both donor 1 and donor 2 experiments (**Fig. 2A**).

**Fig. 2.**
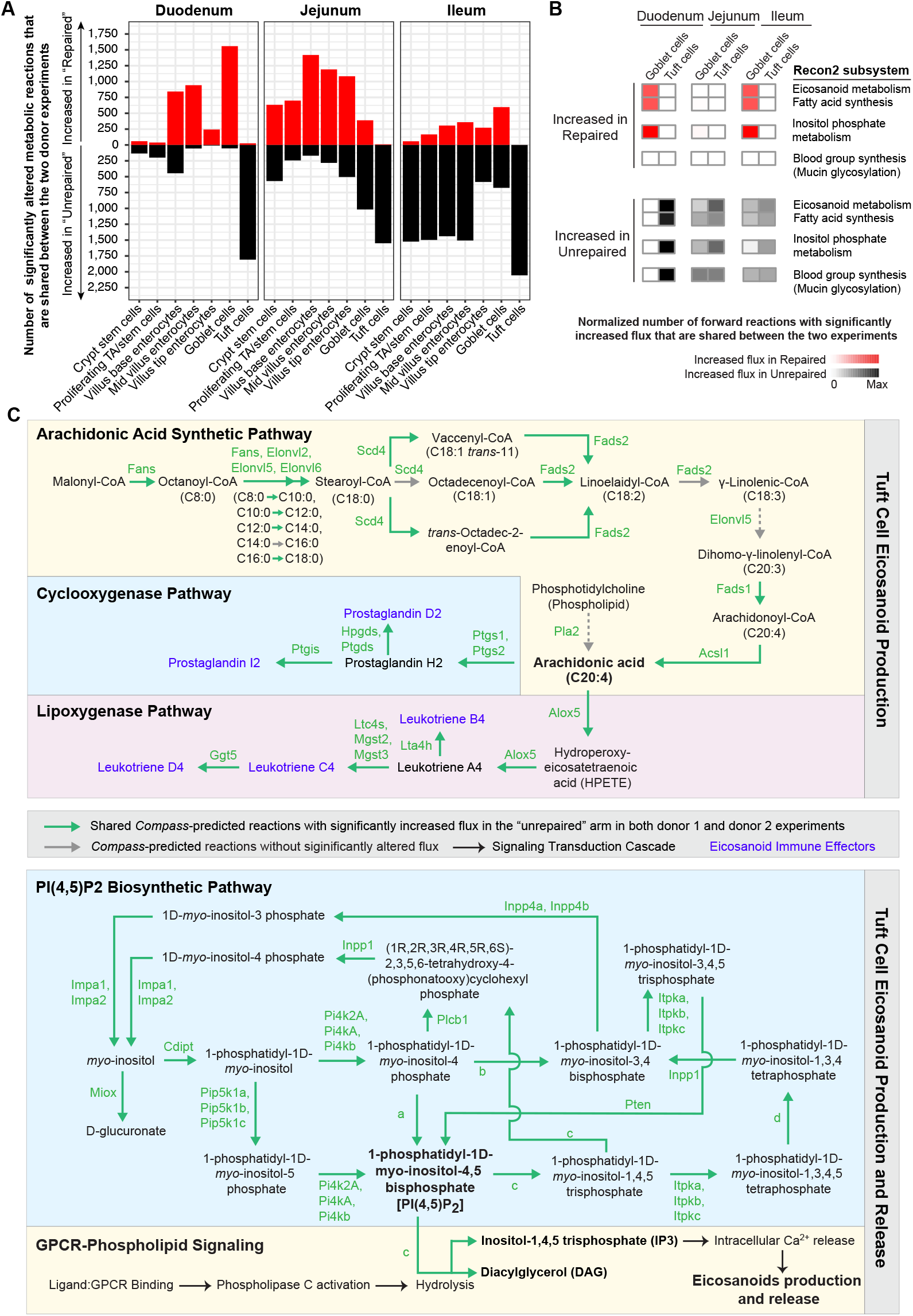
Compass-based analysis of tuft cell metabolism in mice modeling unrepaired and repaired microbiomes. **A**. Bar plot showing the number of metabolic reactions with significantly altered flux, as predicted by *Compass*, shared between donor 1 and donor 2 experiments. Data are presented for each segment of small intestine, including duodenal, jejunal, and ileal regions. **B**. Heatmap representing the normalized number of Compass-predicted forward metabolic reactions with significantly altered flux that are shared between donor 1 and donor 2 experiments. Data are presented for select Recon2 metabolic subsystems in tuft cells and goblet cells from duodenum, jejunum and ileum. The blood group synthesis subsystem includes glycosylation reactions that are also involved in mucin glycosylation, as indicated by the parenthesis. Red indicates the normalized number of reactions with increased flux in the repaired arm while black indicates the normalized number of reactions with increased flux in the unrepaired arm. Color density represents the number of reactions with significantly increased flux in a given Recon2 subsystem within a specific cell type of an experimental arm, normalized to the total number of such reactions across all epithelial cell clusters in the same arm. **C**. Top panel is a schematic of tuft cell metabolic pathways predicted by Compass to exhibit significantly altered flux in response to unrepaired and repaired microbiomes. These pathways include arachidonic acid synthesis, prostaglandin synthesis (via the cyclooxygenase pathway), and leukotriene synthesis (via the lipoxygenase pathway). Bottom panel is a schematic representation of metabolic pathways involved in PIP2 synthesis and GPCR-phospholipid signaling in tuft cells, which facilitate the release of eicosanoid immune effectors. Enzyme group a includes Pikfyve, Pip4k2a, Pip4K2b, Pip4K2c, Pip5k1a, Pip5k1b, Pip5k1c. Enzyme group b includes Hcst, Pik3c2a, Pik3c2b, Pik3c2g, Pik3ca, Pik3cb, Pik3cd, Pik3cg, Pik3r1, Pik3r2, Pik3r3, Pik3r5. Enzymes group c includes Plcb1, Plcb2, Plcb3, Plcb4, Plcd1, Plcd3, Plcd4, Plce1, Plcg1, Plcg2, Plch1, Plch2, Plcl1, Plcxd2, Plcz1. Enzyme group d includes Inpp5a, Inpp5b, Inpp5d, Inpp5e, Inpp5j, Inppl1, Synj1. Green arrows indicate shared reactions with significantly increased flux in duodenal, jejunal, and ileal tuft cells of the “unrepaired” arm in both donor 1 and donor 2 experiments. Grey arrows represent reactions without significant flux changes. Black arrows denote the signal transduction cascade associated with GPCR-phospholipid signaling. Blue text highlights the eicosanoid immune effectors synthesized in tuft cells. Green texts represent enzymes involved in catalyzing the metabolic reactions.

Across all three regions of the small intestine, multiple epithelial cell types exhibited high degree of overlap in their Compass-predicted metabolic changes across the two donor experiments (**Dataset S3C**). In mice that received repaired microbiomes, crypt stem cells, proliferating TA/stem cells, and the three enterocyte clusters had transcriptional changes indicative of increased activities of pathways related to glutamate metabolism and fatty acid utilization - two key sources of energy for these cells (11–13) (**SI Appendix Fig. S2A; Dataset S3D**).

#### Tuft cell response

Tuft cells play a key role in modulating mucosal immunity, in part by secreting eicosanoid immune effectors including leukotrienes and prostaglandin D_2_ (14). *Compass* analysis revealed that tuft cells from mice colonized with unrepaired microbiomes exhibited increased flux through multiple metabolic pathways involved in the biosynthesis and release of these effectors; they include elevated activities in the fatty acid metabolism, eicosanoid metabolism and inositol phosphate metabolism subsystems (**Fig. 2B**). Specifically, flux in reactions leading to the synthesis of arachidonic acid, a common precursor for synthesis of leukotrienes and prostaglandins, was increased (**Fig. 2C**). Downstream of arachidonic acid, reactions leading to the synthesis of prostaglandins and leukotrienes were predicted to have significantly increased flux (**Fig. 2C**). Release of these eicosanoid effectors depends on an upstream GPCR signaling cascade that involves a critical metabolic reaction in which 1-phosphatidyl-1D-*myo*-inositol-4,5-bisphosphate (PI(4,5)P_2_) is hydrolyzed to second-messengers inositol-1,4,5 triphosphate (IP_3_) and diacylglycerol (DAG) (15). IP_3_ binds to intracellular calcium channels in the endoplasmic reticulum, causing release of Ca^2+^ into the cytoplasm, which in turn triggers release of eicosanoid immune effectors. The snRNA-Seq data-derived *Compass*-based analysis of tuft cells indicated enhanced flux towards biosynthesis of PI(4,5)P_2_, the precursor for IP_3_, in the unrepaired donor 1 and donor 2 arms across all three small intestinal segments (**Fig. 2C**).

We used targeted ultrahigh performance liquid chromatography-triple quadrupole mass spectrometry (see *Method*s) to assay prostaglandins (D_2_, E_2_, I_2_) and leukotrienes (B_4_, C_4_, D_4_) in duodenal, jejunal and ileal segments (n=4 mice/arm; 4 arms). However, all these compounds were below the limit of detection (< 45ng/mg tissue), likely reflecting the fact that tuft cells are sparsely represented (0.4 to 2% of cells) in the small intestinal epithelium (16).

Activation of G protein coupled receptor (GPCR) signaling in tuft cells can form a positive feedback loop to promote tuft cell differentiation and hyperplasia from intestinal stem cells (17–19). Using antibodies to Doublecortin-Like Kinase 1 (Dclk1), we quantified the number of tuft cells in sections prepared from the duodenum, jejunum and ileum (n = 4 mice/treatment group; 4 treatment groups). The number of tuft cells was significantly increased in the duodenum and jejunum of mice colonized with the unrepaired compared to repaired microbiome from donor 1 (Mann-Whitney U Test; P=0.015 and 0.030, respectively), with a trend towards increased numbers in the colon (*P=*0.056) (**Fig. 3A; Dataset S4**). Tuft cell number was significantly increased in the jejunum (*P*=0.015), ileum (*P*=0.015) and colon (*P*=0.015) in mice harboring the unrepaired microbiome from donor 2, with a trend towards an increase in the duodenum (*P*=0.056) (**Fig. 3A; Dataset S4**).

**Fig. 3.**
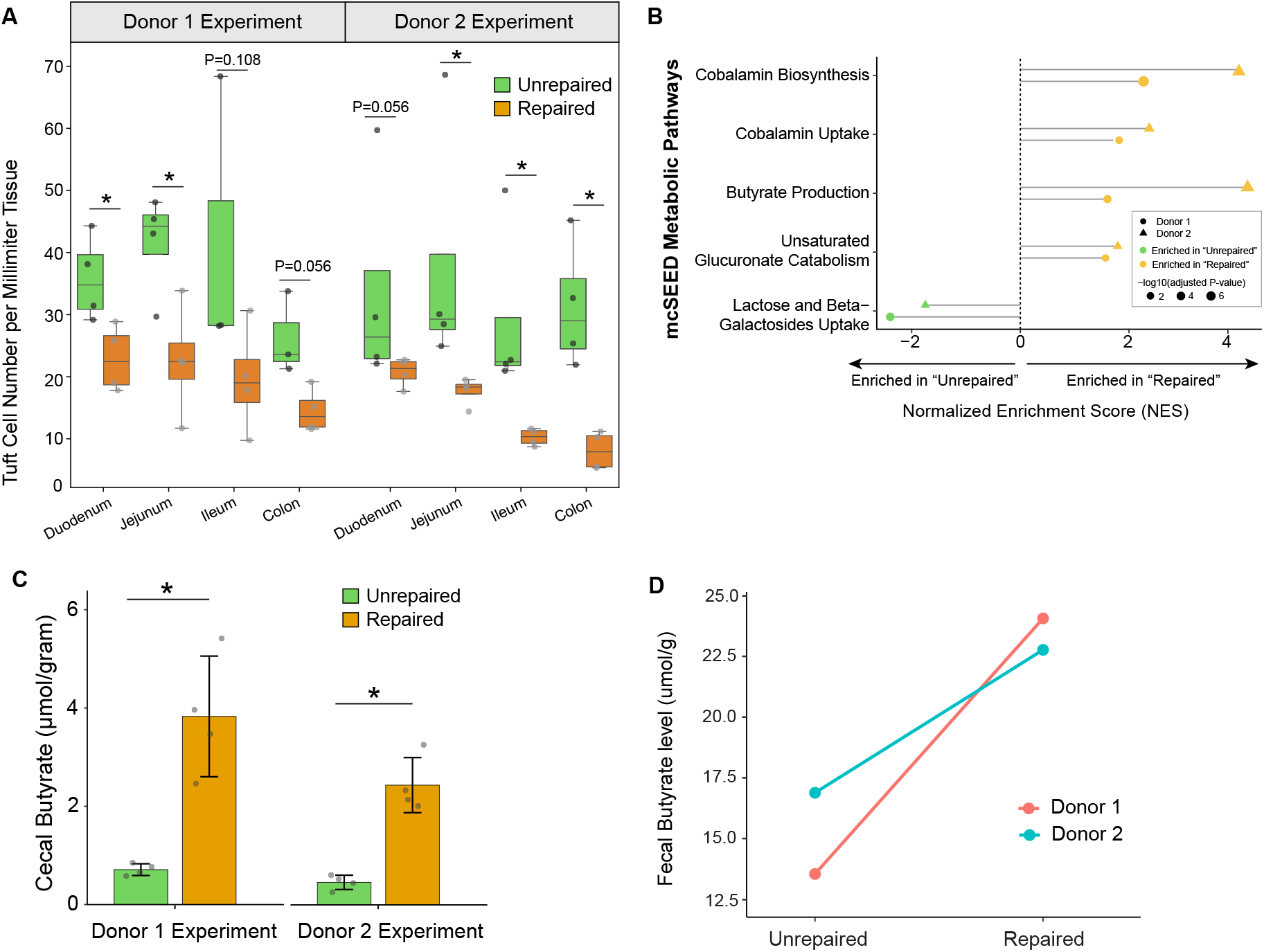
Tuft cell hyperplasia and cecal butyrate level changes in response to “unrepaired” gut luminal ecosystem. **A**. Quantification of tuft cell density across different intestinal regions (duodenum, jejunum, and ileum) based on whole-slide scanned images of DCLK1-immunostained sections. Data are presented as whisker box plots for each donor and intestinal segment. **\***, *P* < 0.05 (Mann-Whitney U test). **B**. Lollipop plot illustrating the normalized enrichment score (NES) for mcSEED metabolic pathways with significantly altered expression from microbial RNA-seq data, as identified from GSEA. The length of each lollipop represents the NES, the color indicates the arm (unrepaired” or repaired) in which the pathway is enriched, and the size of the dots reflects the level of statistical significance of the enrichment (adjusted P-value). **C**. Cecal butyrate levels (μmol/g cecal contents). **D**. Fecal butyrate levels (μmol/g feces) in samples obtained from donor 1 and donor 2 prior to MDCF-2 treatment (unrepaired) and at the conclusion of the 3-month intervention (repaired).

#### Expression of metabolic pathways in the transplanted microbiomes

We performed an *in silico* analysis to define the representation of 158 metabolic pathways described in microbial community SEED (mcSEED) in all MAGs (**Dataset S5**) (20). We used the findings to interpret the results of microbial RNA sequencing of cecal contents collected from pups on P32 (n=4 animals per arm; 2 arms per donor experiment; **Dataset S5A**,**B**). Gene Set Enrichment Analysis (GSEA) was then applied to identify mcSEED pathways whose expression was significantly enriched (*q*-value < 0.1) in the MAG-derived meta-transcriptomes. We focused on pathways that were significantly altered by microbiome repair in both transplanted donor 1 and donor 2 communities (**Dataset S5C)**.

Transcripts assigned to the ‘lactose and beta-galactosides uptake’ pathway were enriched in the metatranscriptomes of the transplanted *unrepaired* donor microbiomes (**Dataset S5C**). Consistent with their known role in lactose utilization, MAGs assigned to *Bifidobacterium longum* and *Bifidobacterium breve* were primary contributors of this pathway, accounting for 10% and 20% of the leading-edge transcripts in the donor 1 experiment, and 12.5% and 37.5% in donor 2 experiment, respectively (**Dataset S5D**). These *Bifidobacterium* species are typically early colonizers of the infant gut, with their representation normally diminishing during the transition from exclusive milk feeding to a weaning diet (21–24). All MAGs assigned to *B. longum* and *B. breve* showed significant *negative* correlations with WLZ in the clinical study performed in 12-18-month-old children with MAM (**Dataset S5D**).

In contrast, both repaired microbiomes were enriched for transcripts involved in (i) butyrate fermentation, (ii) unsaturated glucuronate catabolism and (iii) cobalamin biosynthesis and uptake (**Fig. 3B**; **Dataset S5C**). Butyrate serves as an important energy source for enterocytes (25) and has anti-inflammatory effects including suppression of NF-κB signaling (26), restriction of tuft cell differentiation and type II immunity (27) and enhancement of gut barrier function (28). The glucuronate catabolic pathway breaks down glucuronic acid to UDP-xylose, a sugar donor required for synthesis of glycosaminoglycans (GAGs) (29–31). In bacteria, GAGs play roles in adhesion and colonization of the gut while in the host, GAGs are integral components of the mucosal barrier (32). Cobalamin and its component corrinoids have important roles in bacterial metabolism and communication, impacting microbial-microbial and microbial-host interactions (33). MAGs assigned to species that contributed to each of these enriched pathway’s leading-edge transcripts are described in **Dataset S5D** which also includes information about whether the MAGs were significantly positively or negatively associated with WLZ in the clinical trials.

Butyrate is a key suppressor of tuft cell activation (27). Consistent with this finding, cecal butyrate concentrations, quantified by gas chromatography-mass spectrometry, were significantly elevated in the repaired compared to unrepaired community contexts for both donors (**Fig. 3C**), while other short-chain fatty acids and related metabolites, such as succinate, were not affected (**Dataset S6**). Fecal butyrate levels in both donors were also increased at the end of MDCF-2 treatment (**Fig. 3D**). Our previous report (6) showed that the butyrate production pathway was significantly enriched in MAGs with high WLZ-association coefficients, further underscoring the clinical relevance of this finding.

#### Goblet cell response

Prostaglandin D_2_ produced by tuft cells is known to act on goblet cells to boost their mucus production and secretion (34, 35). Immunostaining of MUC2 disclosed a significant increase in goblet cell density along the length of the small intestines of mice colonized with unrepaired microbiomes from donors 1 and 2 (**Fig. 4B**; **Dataset S7**; n = 1 section per intestinal region, 4 mice per arm; 2 arms per donor experiment).

**Fig. 4.**
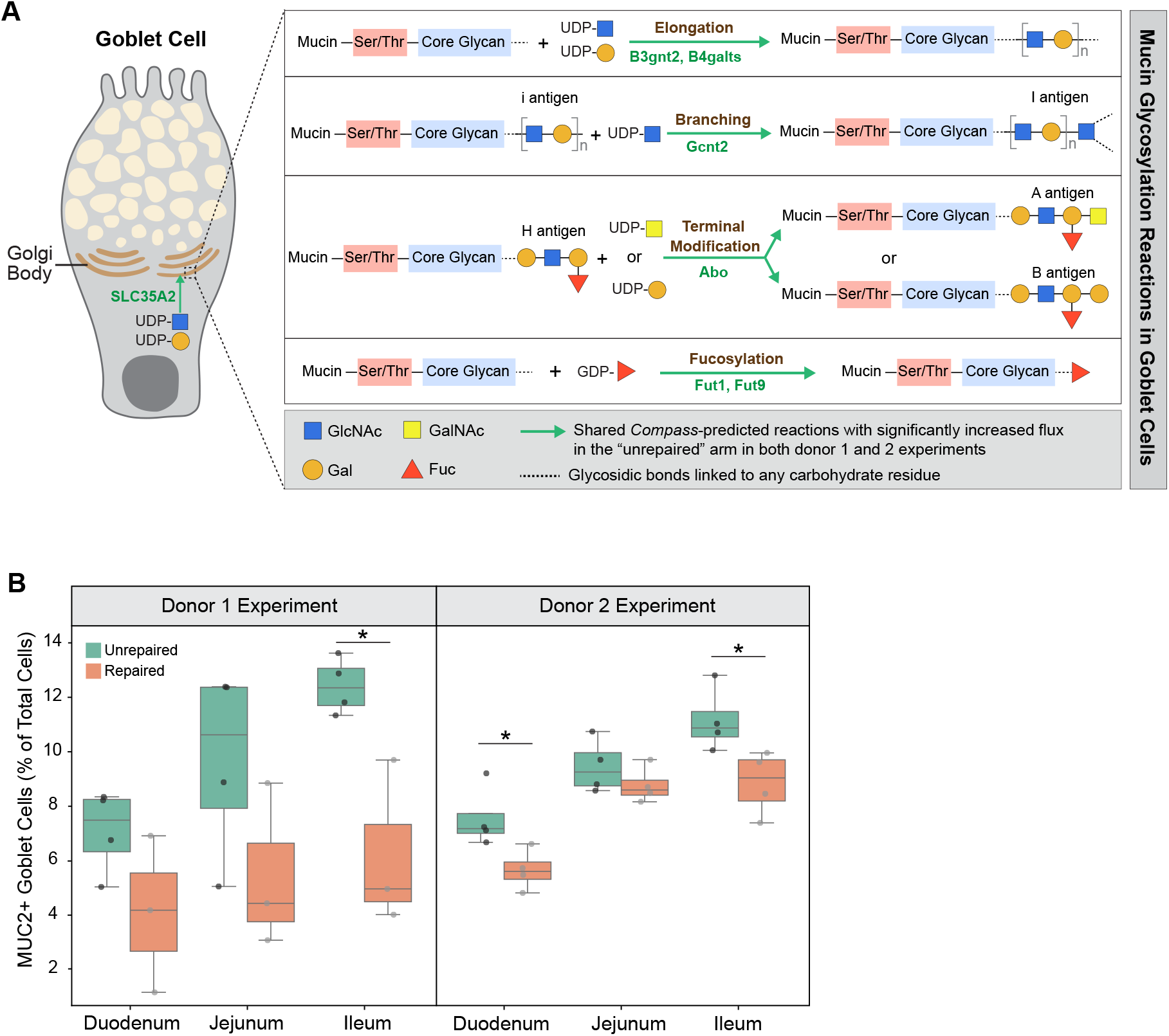
Goblet cell hyperplasia and mucin glycosylation in response to “unrepaired” gut luminal ecosystem. **A**. Schematic representation of four primary types of mucin glycosylation reactions predicted by *Compass* to have significantly increased flux in goblet cells exposed to the “unrepaired” microbiome. These include glycan elongation, branching, terminal modification, and fucosylation. Abbreviations: GlcNAc, N-acetylglucosamine; GalNAc, N-acetylgalactosamine; Gal, Galactose; Fuc, Fucose; Ser/Thr, Serine/Threonine. Green arrows indicate shared reactions with significantly increased flux in jejunal and ileal goblet cells present in the “unrepaired” arm across both donor 1 and donor 2 experiments. Grey arrows represent reactions with no significant changes in flux. Grey dotted lines denote glycosidic bonds linked to any carbohydrate residue. Green text represents enzymes involved in catalyzing the metabolic reactions. **B**. Goblet cell density across three intestinal regions (duodenum, jejunum, and ileum) based on whole slide scanned images of MUC2 immunostaining. Data are presented as whisker box plots for each donor and intestinal segment. **\***, *P* < 0.05 (Mann-Whitney U test).

Mucin, the principal component of the mucus layer, undergoes extensive, multi-step glycosylation in the Golgi apparatus of goblet cells before secretion; these steps are critically linked to its barrier-protecting properties (36, 37). Compared to mice colonized with repaired microbiomes, jejunal and ileal goblet cells from mice harboring unrepaired communities exhibited significantly greater enzymatic flux in the Recon 2 subsystem annotated as “blood group biosynthesis”; this subsystem encompasses the same glycosylation reactions used in the production of mucin (**Fig. 2B**). The first reaction involves enhanced mucin glycan chain elongation mediated by β-1,4-galactosyltransferases (B4GALT) and β-1,3-N-acetylglucosaminyltransferase 2 (B3GNT2). B4GALT incorporates galactose (Gal) residues, while B3GNT2 adds N-acetylglucosamine (GlcNAc) residues, together forming repeating disaccharide units [- Galβ(1,4)-GlcNAcβ(1,3)-] (37). *Compass* also predicted enhanced transport of the two essential nucleotide-sugar donors, UDP-Gal and UDP-GlcNAc into goblet cell Golgi compartments via the transporter SLC35A2 (**Fig. 4A**). The second type of reaction identified by *Compass* as increased in the goblet cells of mice colonized with the unrepaired microbiomes involves N-acetylglucosaminyltransferases (GCNTs), which facilitate glycan branching by adding N-acetylglucosamine (GlcNAc) residues to existing i-antigen structures in mucin glycans (**Fig. 4A**). The third category of glycosylation reactions predicted to be increased in mice that modeled the unrepaired microbiome state is related to the formation of A and B antigen structures as a means of terminal modifications on mucin glycan chains. Catalyzed by A and B transferases encoded by the Abo gene, N-acetylgalactosamine (GalNAc) and galactose (Gal) residues are transferred to the H antigen, respectively. The presence of these ABH antigens in glycan termini influences mucosal interactions with microbes (36). The fourth type of reaction involved fucosylation, catalyzed by members of the fucosyltransferase (FUT) family. Despite being a minor component of mucin glycans, fucose is vital for enhancing mucus viscoelasticity and forming key terminal mucin glycan structures (36).

Goblet cells rely on GPCR-phospholipid signaling to trigger mucus secretion. In mice that received both unrepaired microbiomes, jejunal goblet cells exhibited increased metabolic flux toward the production of IP3 and DAG (**SI Appendix Fig. S3**). IP3 stimulates the release of Ca^2^□ from the endoplasmic reticulum, a signaling event that is crucial for mucus secretion from goblet cells (15, 38).

Expansion of tuft and goblet cells in the gut epithelium is commonly associated with host defense mechanisms triggered by enteric parasites (17–19). To exclude parasite infection as a contributing factor, we aligned shotgun sequencing reads from cecal contents harvested from P32 mice to EuPathDB (release 48) (39), a curated genomic database containing 388 eukaryotic pathogens that includes intestinal parasites such as protists that are capable of inducing tuft cell responses (17–19). Metagenomic reads were pre-filtered to remove host and bacterial MAG sequences prior to alignment. None of the samples contained detectable levels of the eukaryotic pathogens represented in this database (see *Methods;* n = 4 animals treatment arm; 4 arms in total; 16 samples). As noted above, microbial RNA-seq analysis revealed significant upregulation of butyrate production pathways in bacterial MAGs from the repaired microbiomes (**Fig. 3C; Dataset S5C**). Together, these findings support the notion that shifts in bacterial metabolism, rather than parasitic infection, underlie the tuft cell response observed in our gnotobiotic model.

### Effects on expression of virulence factors and components of epithelial junctions

We used DIAMOND (40) to align the protein sequences of virulence factors from the Virulence Factor Database (VFDB) to the protein-coding sequences of MAGs in cecal and fecal samples from postpartum day 32 dams and their P32 pups that satisfied colonization criteria for abundance and prevalence (see *Methods*). A MAG was deemed ‘putatively virulent’ if it encoded both an exotoxin and an effector delivery system component (DIAMOND alignment criteria: bitscore >200 and E-value <10^-100^; **Dataset S2E**). The results disclosed that samples collected from animals with either of the two donors’ unrepaired microbiomes contained a significantly greater number of MAGs satisfying these virulence factor criteria than the corresponding repaired microbiomes (**SI Appendix Fig. S4A**). GSEA of the microbial RNA-Seq datasets revealed significantly enriched expression of VFDB pathways involved in (i) host extracellular matrix degradative functions, (ii) effector delivery systems (injection of virulence factors into host cells), (iii) immune modulation (host immune system evasion, (iv) stress survival and (v) a Salmonella pathogenicity island in cecal samples obtained from mice harboring the unrepaired microbiomes (**Fig. 5A; Dataset S5E**,**F**). Biofilm formation was the only VFDB pathway enriched in a repaired microbiome (**Fig. 5A; Dataset S5E**,**F**). Although biofilms support bacterial persistence, they are not necessarily indicative of a state of increased virulence.

**Fig. 5.**
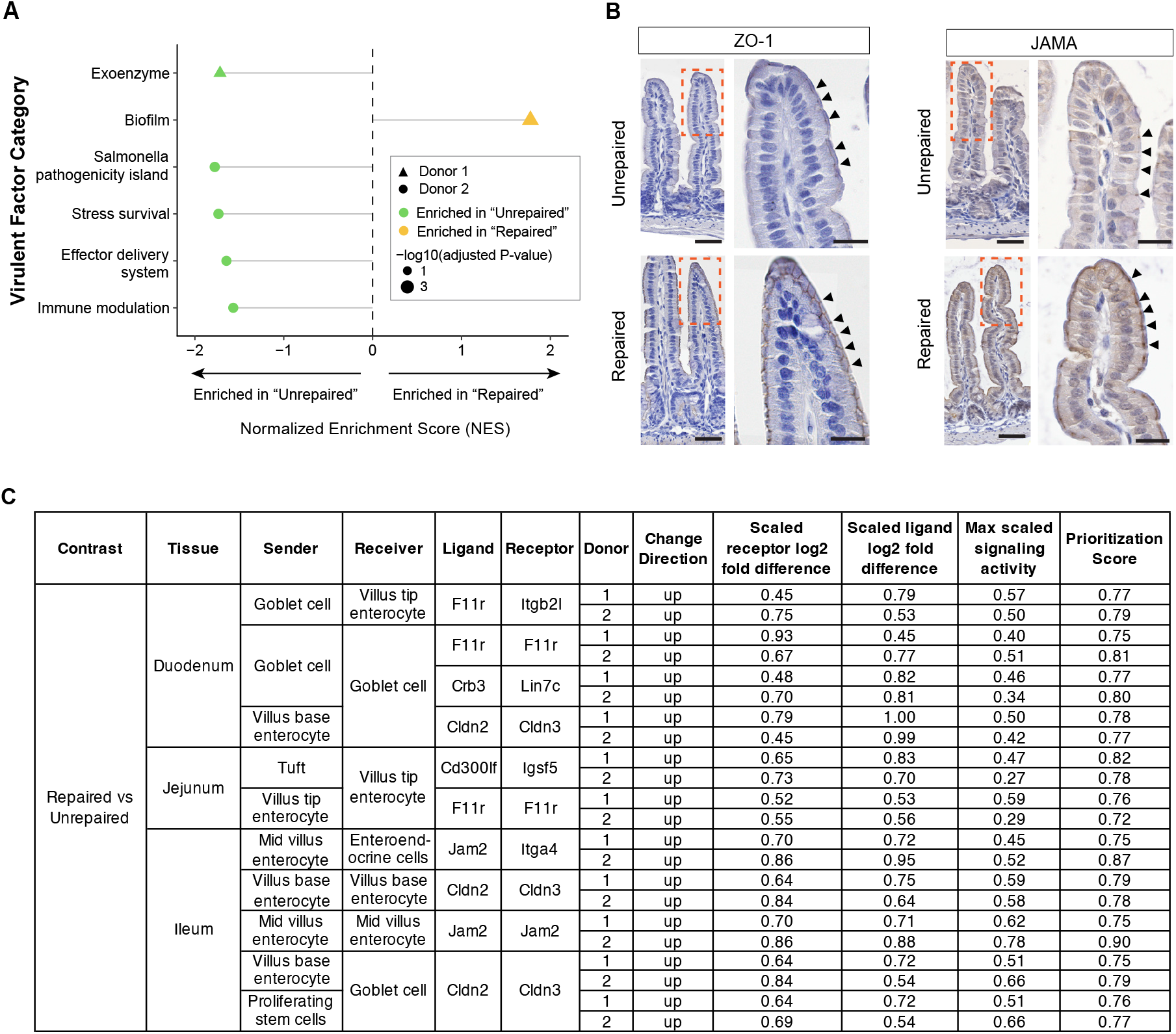
Alterations in microbial virulence and epithelial cell tight junctions. **A**. Lollipop plot of NES for virulence factor categories from VFDB with significant altered expression, as identified by GSEA from microbial RNA-seq data. **B**. Representative immunostaining images showing the distribution of tight junction components ZO-1 and JAMA in the jejunums of P32 mice. Brown staining indicates positive immunoreactive signal, while blue staining is from nuclei counterstained with hematoxylin. Scale bars: 50 μm in the left panels and 20 μm the right panels. **C**. Table summarizing the shared epithelial tight junctional interactions that ranked in the top 10% prioritization score across duodenum, jejunum and ileum in mice from both donor 1 and donor 2 experiments.

Immunostaining for epithelial tight junction biomarkers ZO-1 and JAMA (encoded by F11r gene) showed decreased levels of both proteins in the duodenums, jejunums and ileums of mice colonized with the two unrepaired donor microbiomes (**Fig. 5B**). In contrast, levels of both proteins were comparable in jejunal tissues from P32 germ-free mice fed weaning diets supplemented with either Mirpur-18 or MDCF-2, indicating that these junctional changes were not driven by diet alone (**SI Appendix Fig. S4B**).

We then applied the “MultiNicheNet” algorithm to the snRNA-Seq dataset to further examine the correlation between microbiome repair and expression of tight junction components that could mediate interactions between various epithelial cell types along the crypt-villus and duodenal to ileal axes of the small intestine (**Dataset S3E**). “MultiNicheNet” infers differentially active ligand-receptor interactions between experimental conditions. For every ligand-receptor pair within a sender–receiver cell type combination, a prioritization score is computed based on parameters described in *Methods*. The higher the prioritization score, the more confidently the model predicts that this ligand-receptor pair is altered between experimental arms. For our analysis, we tested all possible epithelial cell-to-cell interactions by designating every epithelial cell type as both a sender and a receiver; we focused on ligand-receptor interactions that ranked in the top 10% prioritization score in both donor 1 and donor 2 experiments. The results are summarized in **Fig. 5C** and indicate that repair is accompanied by increased expression of junctional components, including JAMs and Claudins, operating between enterocytes themselves, between enterocytes and tuft cells, goblet cells and enteroendocrine cells, as well as between goblet cells themselves and goblet and transit amplifying cells. Remarkably, the effects of microbiome repair on expression of junctional components varies as a function of cellular position along the crypt-to-villus axis (*e*.*g*. Cldn2, expressed in enterocytes located at the base of villi) as well as the duodenal-ileal axis (JAMA, restricted to the duodenum and jejunum). Collectively, these results provide evidence supporting a relationship between microbiome repair and fortification of the gut epithelial barrier.

## DISCUSSION

In the present study, we employed a ‘reverse translation’ approach that used gnotobiotic mice colonized with intact uncultured fecal samples from two Bangladeshi children with moderate acute malnutrition (MAM) enrolled in a completed, randomized controlled trial of a microbiome-directed therapeutic food. The experimental design featured an ‘unrepaired arm’ where a group of gnotobiotic pups received, from their dams, a fecal microbiome sample obtained from a trial participant just before initiation of MDCF-2 treatment, and a ‘repaired arm’ where a corresponding group of pups received a microbiome sample obtained from the same child on the last day of treatment. The cellular and molecular effects of the unrepaired and repaired communities from the two children with MAM on gut biology was examined the context of diets representative of those consumed by the children at the time of their microbiome sampling. The two human donors used in this preclinical study were selected based on their robust ponderal growth responses (upper quartile of trial participants receiving MDCF-2), and the accompanying robust change in the representation of growth-associated bacterial taxa.

Compared to mice receiving repaired microbiomes, animals that received the unrepaired pretreatment communities exhibited features of a more virulent gut luminal ecosystem that were reflected in the responses of intestinal tuft cells and goblet cells. These features included (i) increased expression of virulence factors by bacterial members of the microbiome, (ii) reduced production of butyrate, a key suppressor of tuft cell activation by the microbiome, (iii) increased number of tuft cells and accompanying increases in their synthesis and metabolism of eicosanoid immune effectors, plus (iv) increased expression of multiple steps in mucin glycosylation in goblet cells. Tuft cells are known to play an important role in type 2 immunity, particularly in response to parasitic infections (17–19). Upon sensing parasites, they secrete IL-25 (17–19), leukotrienes (41–43) and prostaglandin D_2_ (44), which in turn activate group 2 innate lymphoid cells (ILC2s). Activation leads to secretion of type 2 cytokines IL-4/13, which act on goblet cells to promote goblet cell hyperplasia (17) and glycosylation of mucin (45). In addition, Prostaglandin D_2_ produced by activated tuft cells directly enhances goblet cell mucus production (34, 35). Chronic activation of tuft cell–mediated immunity, as observed in parasitic infection models, has been linked to moderate weight loss (14), suggesting a potential mechanism underlying the reduced weight phenotype observed in our experiments.

To date, only a limited number of metabolites, including butyrate, succinate and N-decanoylglycine, have been shown to modulate tuft cell number/ biology. Our study documented an increase in levels of butyrate, a known inhibitor of tuft cell expansion, under microbiome repaired compared to unrepaired conditions. There were no significant differences in levels of succinate, a well-characterized metabolite that activates tuft cells. N-decanoylglycine was recently identified as a tuft cell– expanding metabolite in a *Shigella* infection model (35) and it was below the limits of detection by LC-QqQ-MS. A small number of studies have begun to implicate tuft cell biology in the context of malnutrition, but significant gaps remain. For example, tuft cell numbers were markedly increased in marmosets with Marmoset Wasting Syndrome (46). Moreover, in gnotobiotic mice colonized with a consortium of bacterial strains cultured from the fecal microbiota of a stunted Malawian infant and fed a representative Malawian infant diet supplemented with sialylated milk oligosaccharides that were (i) increases in intestinal succinate levels and in the number of tuft cells, (ii) activation of a succinate-induced pathway in tuft cells linked to Th2 responses and (iii) reductions in osteoclastogenic activity and bone resorption. These effects were not observed in germ-free animals fed the same supplemented diet, highlighting the critical role played by the microbiota in mediating this tuft cell phenotype (47).

We chose to use a gnotobiotic model that involved maternal-to-pup microbiome transmission rather than one that that employed direct introduction of microbial communities into already weaned animals. We did so because early-life colonization is a critical window for intestinal and immune maturation. The postnatal time points selected for analyzing offspring (P14 and P32) correspond to the suckling and post-weaning periods in humans, providing a window for modeling how MDCF-2– associated microbiome repair influences gut function and growth.

Our model can be used to extend these analyses in the future. Integrating the results of microbial RNA-seq with non-targeted and targeted mass spectrometry provides an opportunity to (i) nominate candidate effectors and biomarkers of tuft cell responses to microbiome repair and (ii) further ascertain the contributions of tuft cells to various facets of intestinal responses/adaptations to undernourished states (including gut mucosal barrier function). The efficiency of mother-to-offspring transmission of MAGs whose abundances were significantly associated with ponderal growth (WLZ) in the clinical trial was high, ranging from 86% to 98%. In principle, the experimental approach could be generalized and be either retrospective or prospective. For example, microbiome samples could be obtained from a given trial participant or combined from multiple trial participants with established robust versus weak responses to treatment. The microbiomes could also be from children representing different ages or different time points during and following treatment, or with different types/degrees of co-morbidities. Moreover, sampled microbiomes could be utilized to compare responses to additional treatments not incorporated into the original trial design or to compare responses of trial participants to those obtained from other untreated populations.

The current study deliberately deferred testing the combination of unrepaired intact community and MDCF-2, or repaired intact community and pre-treatment diets, largely because directly determining the effects of food components and community membership on the representation of expressed microbial and host functions requires an ability to deliberately and precisely manipulate these two factors. Given that microbial communities adapt rapidly to dietary changes (48), introducing a ‘mismatch’ between a microbiome’s repair status and diet (i.e., combining an unrepaired microbiome with a MDCF-2-supplemented weaning diet or a repaired microbiome with an unsupplemented weaning diet) could yield ‘hybrid’ communities that do not reflect the states in clinical trial participants that were the focus of our mechanistic analyses. A previous study from our group that used a simplified microbiota comprised of a consortium of cultured genome-sequenced age- and growth-discriminatory organisms cultured from the study population did not detect the effects of MDCF-2 treatment on tuft and goblet cells (8). However, integrating findings from reverse translation experiments employing intact microbiomes with experiments involving defined consortia should enable further refinement of the composition of defined communities and tests of hypotheses about the mechanisms by which microbiome repair affects microbial community and host biology. For example, adding enteropathogens cultured from trial participants to the defined community of cultured age- and growth-discriminatory bacterial taxa that we previously examined in gnotobiotic mice (8), represents one approach for deciphering the mechanisms by which the MDCF-2 and/or future prebiotic/synbiotic formulations designed to effect microbiome repair and promote growth can result in pathogen exclusion and affect components of the mucosal barrier.

## METHODS

### Gnotobiotic mouse studies

Gnotobiotic mouse experiments were conducted using protocols approved by the Washington University Animal Studies Committee. Germ-free C57BL/6J mice were housed in plastic flexible film isolators (Class Biologically Clean Ltd) maintained at 23 ºC with a strict 12-hour light/dark cycle (lights on at 0600h). To support natural nesting behaviors and provide enrichment, autoclaved paper ‘shepherd shacks’ were included in each cage. Weaning diets supplemented with Mirpur-18 and MDCF-2 were prepared as described previously (8) and sterilized via gamma irradiation (30–50 KGy). The sterility of the pellets was confirmed by culturing them in LYBHI medium and Wilkins-Chalgren Anaerobe Broth under aerobic and anaerobic conditions for 7 days at 37 ºC, followed by plating on LYBHI and blood-agar plates. Nutritional analysis was conducted by Nestlé Purina Analytical Laboratories (St. Louis, MO) (**Dataset S1**).

### Donor selection and preparation of fecal microbial community samples

The study used fecal samples collected prior to treatment (day 0) and end-of-treatment (day 90) from two participants (see **Dataset S2C** for donor characteristics) in the MDCF-2 treatment arm of a previously reported randomized controlled clinical trial (ClinicalTrials.gov identifier NCT04015999) that was conducted in Dhaka, Bangladesh and approved by the Ethical Review Committee at the icddr,b (5).

Donors were selected based on the following criteria: (i) their WLZ improvement ranked within the upper quartile among all participants in the MDCF-2 arm; (ii) they exhibited the greatest number and largest magnitude of increases in bacterial taxa positively associated with WLZ in response to MDCF-2 treatment; and (iii) they showed the most extensive reductions in both number and magnitude of taxa negatively associated with WLZ (8).

To prepare intact, uncultured fecal microbial communities, fecal samples were pulverized in liquid nitrogen. Aliquots of pulverized material (300 mg) were transferred to a Coy Chamber under anaerobic conditions (75% N_2_, 20% CO_2_ and 5% H_2_) and resuspended in 4 mL of phosphate buffered saline (PBS) containing 30% glycerol and 0.05% cysteine-HCl. The mixture was vortexed for 2 minutes, pausing for 5 seconds every 30 seconds, and passed through a 100 µm pore-diameter strainer (Corning, catalog number 431752). One milliliter aliquots of the resulting filtrate were placed into crimp-top tubes (Wheaton, catalog number 225175) which were transferred out of the Coy Chamber and stored at -80 ºC until use. A 200 µL aliquot was administered to dams via a 22-gauge oral gavage needle (Cadence Science, catalog number 7901).

### Metagenomic sequencing

DNA was isolated from cecal contents and fecal samples by bead beating and phenol-chloroform extraction and further purified using the QIAquick 96 PCR Purification Kit. Shotgun sequencing libraries were prepared using the Nextera XT DNA Library Prep Kit and sequenced using an Illumina NextSeq platform (2x150 bp paired-end reads). Sequencing yielded an average of 6.15 × 10□ ± 3.3 × 10□ reads per sample from the donor 1 experiment and 6.04 × 10□ ± 2.24 × 10□ reads per sample from the donor 2 experiment (mean ± SD; **Dataset S2A**). Reads were demultiplexed using bcl2fastq (v2.2.0), trimmed with Trim Galore (v0.6.4), and filtered to exclude host-derived sequences using bowtie2 (v2.3.4.1). The remaining reads were aligned using Kallisto (v0.43.0) to the 1000 MAGs that had previously been assembled from participants in the randomized controlled clinical trial (6) to generate a transcript-per-million (TPM) count matrix. Differential abundance analysis between treatment conditions was conducted using DESeq2 (50). MAGs were considered to have colonized the mouse gut if their transcript-per-million (TPM) abundance exceeded 20 and their prevalence across samples was greater than 40%.

To identify putative virulent MAGs, we employed DIAMOND (v2.1.4) (40) to align protein sequences of virulence factors from the Virulence Factor Database (VFDB) (50) against the protein-coding sequences of MAGs identified in P32 dams and pups from “unrepaired” and “repaired” cecal and fecal samples. MAGs were included in the analysis if they met the colonization criteria of TPM >20 and prevalence greater than 40%. A MAG was classified as putatively virulent if it encoded both exotoxins and effector delivery systems, two essential components of bacterial pathogenesis. Alignments were filtered using a bitscore threshold of >200 and an E-value cutoff of <10^-100^.

Reads from shotgun sequencing of cecal DNA samples from P32 pups were filtered to remove host and MAG sequences and then used Kraken2 (51) to align to EuPathDB of 388 eukaryotic pathogens (release 46; https://veupathdb.org) (39).

### Microbial RNA sequencing (microbial RNA-Seq)

RNA was isolated from cecal contents collected from P32 pups using phenol-chloroform based method. cDNA libraries were generated from isolated RNA samples using the ‘Total RNA Prep with Ribo-Zero Plus’ kit (Illumina). Barcoded libraries were sequenced (Illumina NovaSeq instrument; 2x150 bp paired-end reads; n=8 samples for the two arms of experiments involving a given donor; 5.5x10^7^ ± 2.0x10^6^ raw reads/sample (mean±SD) in the donor 1 experiment and 5.4x10^7^ ± 4.7x10^6^ raw reads/sample in donor 2 experiment; **Dataset S3A**]. Raw reads were processed for quality control and trimming by fastp v0.20.0. Trimmed reads were aligned to mouse mm10 genome using bowtie2 v2.4.2 to filter out host reads. Host-filtered reads were mapped to the 1000 MAGs identified in the POC study (6) with Kallisto v0.48.0. The resulting Kallisto pseudocount dataset was imported into R v4.1.4.

GSEA was conducted for (i) virulence factor categories from the Virulence Factor Database (VFDB) and (ii) metabolic pathways annotated in the mcSEED database (20). Using the fgsea package v1.16.0. A ranked gene list was created by ordering genes included in the differential expression analysis in decreasing order based on -log_10_(*P*-value)*sign of log_2_(fold-difference). Gene sets were derived from VFDB categories and from mcSEED module 3. Enrichment analysis was performed with 100,000 permutations, and gene sets with an adjusted P-value of <0.1 were considered significantly enriched.

### Single-nucleus RNA-Sequencing (snRNA-seq) and analysis

The small intestine was divided into thirds; 4 cm long segments was recovered from each third and defined as duodenum, jejunum and ileum. Each 4 cm segment from each mouse was opened longitudinally and incubated in ice-cold PBS containing 10 mM EDTA for 10 minutes. The epithelial layer was lifted by gently brushing and scraping with forceps, collected and snap frozen in liquid nitrogen. Nuclei were isolated from the epithelial layer using a method originally described for brain (52) except that the concentration of bovine serum albumin was increased to 2% and RNase inhibitors (Sigma, catalog number 3335402001) to 0.2U/μL in both the lysis and resuspension buffers. 10,000 nuclei per sample were subjected to gel bead-in-emulsion (GEM) generation, reverse transcription and library construction according to the protocol provided in the 3’ gene expression v3.1 kit manual (10X Genomics). Balanced libraries were sequenced [Illumina NovaSeq S4; 2 x150bp paired-end reads; 6.5±1.6x10^4^ reads/nucleus (mean±SD); **Dataset S4A**].

Read alignment, feature-barcode matrices and quality controls were processed using the CellRanger 5.0 pipeline with the ‘--include-introns’ option to ensure intronic reads would map to the mouse reference genome (GRCm38/mm10). Nuclei with over 2.5% reads from mitochondria-encoded genes reads or ribosomal protein genes were excluded from downstream analysis. Sample integration, count normalization, cell clustering and marker gene identification was performed using Seurat (v4.0). Filtered feature-barcode matrices were converted into Seurat objects using the CreateSeuratObject function (minimum criteria: 5 cells and 200 features). Sample normalization and integration were performed with Seurat (v4.0). The integrated dataset was subjected to unsupervised clustering using the shared nearest-neighbor graph-based algorithms (FindNeighbors and FindClusters; dimensions = 1:30, resolution = 0.8). Cell clusters were manually annotated based on the expression of known marker genes. Differential gene expression across experimental conditions was analyzed based on “pseudobulk” approach as described in ref. 8.

MultiNicheNet v2.0.0 (53) was applied to snRNA-seq datasets generated from the duodenum, jejunum, and ileum in both donor 1 and donor 2 experiments. MultiNicheNet enables inference of condition-specific ligand-receptor interaction changes across defined sender and receiver cell populations. For every ligand-receptor pair within a sender–receiver cell type combination, a prioritization score is computed based on (i) log□-fold change of the ligand expression in sender cells, (ii) log□-fold change of the receptor expression in the receiver cells, (iii) −log□□ *P*-value times the sign of the fold change in the ligand expression in sender cells, (iv) −log□□ *P*-value times the sign of the fold change in the receptor expression in receiver cells, (v) ligand activity score based on the differential expression of the target genes in the receiver cells, (vi) mean expression of the ligand in sender cells, (vii) mean expression of the receptor in receiver cells, and (viii) proportion of samples in which both ligand and receptor are expressed in 10% of sender and receiver cells. The higher the prioritization score, the more confidently the model predicts that this ligand-receptor pair is altered between experimental arms. snRNA-Seq data from seven epithelial cell clusters (crypt stem cells, proliferating transit amplifying cells, villus-base enterocytes, mid-villus enterocytes, villus tip enterocytes, tuft cells, and goblet cells) were subjected to Compass-based *in silico* metabolic flux analysis (version 0.9.10). Compass assigned reaction scores to each Recon2 reaction for each cell cluster. Reaction scores were filtered based on the criteria described in our previous publication (8). Briefly, only Recon2 (54) reactions that are supported by biochemical evidence (defined by Recon2 as having a confidence level of 4) and have complete enzymatic information for the reaction were advanced to the follow-on analysis. This filtering yields 2,075 pass filter reactions in 83 Recon2 subsystems. For each Recon 2 reaction, a Mann-Whitney U test was used to test the statistical significance of the difference in flux between the unrepaired and repaired arm. *P* values from the Mann-Whitney U tests were adjusted for multiple comparisons with the Benjamini–Hochberg method. For reversible reactions, a treatment condition may lead to statistically significant increases or decreases in both the forward and reverse directions within a specific cell cluster. To determine the net directionality in such cases, we compared reaction scores between the two directions. If the score of the forward reaction exceeded that of the reverse reaction, the reaction was classified as net forward; if the converse was true, it was classified as net reverse.

### Histology and Immunostaining

Duodenal, jejunal, and ileal tissues, collected from P32 pups in both donor 1 and donor 2 experiments, were fixed in 10% formalin, embedded in paraffin, and sectioned into 5μm-thick slices along the longitudinal axis of the small intestine. Immunostaining was performed using primary antibodies direct against DCLK1 (Cell Signaling Technology, catalog number 62257S, 1:300 dilution), ZO-1 (Abcam, catalog number ab221546, 1:200 dilution), JAMA (ThermoFisher, catalog number 36-1700, 1:400 dilution) and MUC2 (Thermo Fisher, catalog number PA5-21329, 1:500 dilution), followed by species-appropriate fluorophore-conjugated or HRP-conjugated secondary antibodies and DAB-based color development. Bright field images of whole slides were acquired using a Hamamatsu NanoZoomer 2.0-HT system. Fluorescent whole-slide images were obtained by a 3DHISTECH Panoramic P250 Flash III slide scanner. Tuft cell density in a given small intestinal segment (duodenum, jejunum or ileal) was determined by using a pixel classifier of QuPath v0.5.1 (53). For each intestinal segment from a given animal, we selected the three best preserved and oriented regions, each containing a minimum of 20 crypt–villus units (‘regions of interest’, ROI). The longitudinal length of each tissue segment in the ROI was determined by drawing a smooth line along the muscularis propria. Tuft cell density was calculated by dividing the total number of tuft cells across the three ROIs by the summed tissue length. Goblet cell number (MUC2+) and total cell number (DAPI+) were quantified using the Cellpose Cyto3 model, applied to three ROIs per intestinal region selected using the same criteria as for tuft cell analysis. Goblet cell density in a given small intestinal region was calculated as the total number of MUC2+ cells divided by the total number of DAPI+ cells across the three ROIs. All quantification was performed without double blinding.

### Mass spectrometry

Short-chain fatty acids were quantified by gas chromatography-mass spectroscopy (GC-MS). Aliquots of flash-frozen cecal contents were weighed and supplemented with 10 μL of a mixture of internal standards (20 mM of acetic acid-^13^C_2_,d_4_, propionic acid-d_6_, butyric acid-^13^C_4_, lactic acid-3,3,3-d_3_, and succinic acid-^13^C_4_). After the addition of 20 μL of 33% HCl and 1 mL diethyl ether, the mixture was vortexed vigorously for 10 minutes and centrifuged at 4,000 × g for 5 minutes. The supernatant was transferred to another vial and a second diethyl ether extraction was performed. After combining the two ether extracts, a 60 μL aliquot was removed, combined with 20 μL N-tert-butyldimethylsilyl-N-methyltrifluoroacetamide (MTBSTFA) in a GC auto-sampler vial with a 200 μL glass insert, and incubated for 2 h at room temperature. Derivatized samples (1 μL) were injected with 15:1 split into an Agilent 7890B/5977B GC-MS system.

For measurement of eicosanoids, frozen intestinal tissue segments were weighed in Matrix D tubes (MP Biomedicals; 6913050) followed by addition of 19 volumes of ice-cold methanol (microliters per milligram wet weight of tissue). Samples were homogenized (eight rounds each for 1 minute at room temperature using a BioSpec mini-bead beater with cooling on ice for 1 minute between each round). The resulting homogenates were centrifuged at 12,000 × g for 5 minutes at 4 °C. A 300-μL aliquot of each supernatant was transferred to a new tube and dried in a centrifugal evaporator (LabConco CentriVap). Dried samples were resuspended in 100 μL of 50% methanol and centrifuged at 12,000 × g for 1 minute at 4 °C to ensure that no particulate matter is carried forward to the next step. An 80 μL aliquot of each supernatant was carefully transferred without creating bubbles into an Agilent sample vial (with fixed insert) and 5 μL was injected into a 1290 Infinity II UHPLC system coupled to a 6470 Triple Quadrupole (QqQ) mass spectrometer equipped with a Jet Stream electrospray ionization source (Agilent Technologies).

## Acknowledgements

We thank David O’Donnell and Maria Karlsson for their assistance with mouse husbandry, Martin Meier for his contribution to generating shotgun sequencing and microbial RNA-Seq datasets, Su Deng and Justin Serugo for oversight of biospecimen archives, and Evan Lee, Christopher Suarez, Winne Chen and Carlito Lebrilla for analyzing glycosidic linkages present in the diets administered to gnotobiotic mice.

## Author contributions

Y.W., H.-W.C. and J.I.G. designed the gnotobiotic mouse experiments, which were performed by Y.W. and H.-W.C. I.M and T.A oversaw the clinical trial from which the fecal samples used in this study were collected. Mouse diets were formulated by H.-W.C. with assistance from M.J.B. and were based on the results of human diet surveys provided by T.A. Shotgun sequencing and microbial RNA-Seq datasets were generated by Y.W., and analyzed together with D.M.W., H.M.L., M.C.H. snRNA-Seq datasets were generated by Y.W. and analyzed by Y.W., H.-W.C., and C.K. Y.W., H.-W.C. and J.I.G. wrote this manuscript with assistance from co-authors.

## Data and materials availability

Shotgun sequencing, microbial RNA-Seq and snRNA-Seq datasets have been deposited in NIH Sequence Read Archive (Accession #: PRJNA1262995). Fecal specimens used in these studies were provided to Washington University under a materials transfer agreement with icddr,b.

## SUPPLEMENTARY APPENDIX FIGURES AND LEGENDS

**SI Appendix Fig. S1. Volcano plots for WLZ-associated MAGs colonized in dams and snRNA-seq analysis of the small intestinal tissues. A**. Volcano plots showing the relationship between WLZ-association effect size (β1 coefficient in the linear model shown in Fig. 1) and statistical significance (– log□□ q-value) for all MAGs identified in the clinical trial (gray points). WLZ-associated MAGs that colonized dams are highlighted in orange for each condition: donor 1’s unrepaired and repaired microbiome (top left and right subpanels, respectively); donor 2’s unrepaired and repaired microbiome (bottom left and right subpanels). **B**. UMAP plot of snRNA-seq dataset generate from jejunal tissues in the donor 1 experiment, highlighting clustering of epithelial cell populations. **C**. Dot plot illustrating the expression of known marker genes across cell types identified in the UMAP plot in panel A. The color intensity represents the expression level, while the dot size corresponds to the percentage of cells expressing the marker.

**SI Appendix Fig. S2. Normalized number of Compass-predicted metabolic reactions with significantly altered flux in 79 Recon2 subsystems. A**. Heatmap representing the normalized number of Compass-predicted forward metabolic reactions with significantly altered flux that are shared between donor 1 and donor 2 experiments. Data are presented for 79 Recon2 metabolic subsystems across all epithelial cell clusters from duodenum, jejunum and ileum. Red indicates the normalized number of reactions with increased flux in the repaired arm, while black indicates the normalized number of reactions with increased flux in the unrepaired arm. Color density represents the number of reactions with significantly increased flux in a given Recon2 subsystem within a specific cell type of an experimental arm, normalized to the total number of such reactions across all epithelial cell clusters in the same experimental arm.

**SI Appendix Fig. S3. Metabolic reactions related to glycosylation and secretion of mucin in goblet cells. A**. Schematic representation of reactions involved in PIP2 biosynthesis and GPCR-phospholipid signaling pathways. Green arrows represent reactions with significantly increased flux predicted by *Compass* in both jejunal and ileal goblet cells of the “unrepaired” arm across donor 1 and donor 2 experiments. Green dotted arrows indicate reactions with significantly increased flux in jejunal goblet cells only. Grey arrows denote reactions with no predicted significant changes in flux. Black arrows highlight the signal transduction cascade associated with GPCR-phospholipid signaling.

**SI Appendix Fig. S4. Putative virulent MAG analysis and immunocytochemical analysis of epithelial junction proteins in control germ-free mice. A**. Bar blot demonstrating the number of colonized putative virulent MAGs with differential abundance between unrepaired and repaired arms in the fecal and cecal samples of P32 pups in donor 1 and donor 2 experiments. **B**. Representative images showing the distribution of tight junction components ZO-1 and JAMA in the jejunums of P32 germ-free pups fed with weaning diets supplemented with Mirpur-18 and MDCF-2. Brown staining indicates a positive immunoreactive signal, while blue staining represents nuclei counterstained with hematoxylin. The area denoted with the dashed red box is shown at higher power in the adjacent image. Scale bars: 50 μm in the left panels and 20 μm the right panels.

