## Supplementary figures and images for "Using gnotobiotic mice to decipher effects of gut microbiome repair in undernourished children on tuft and goblet cell function"

### Supplementary Figures

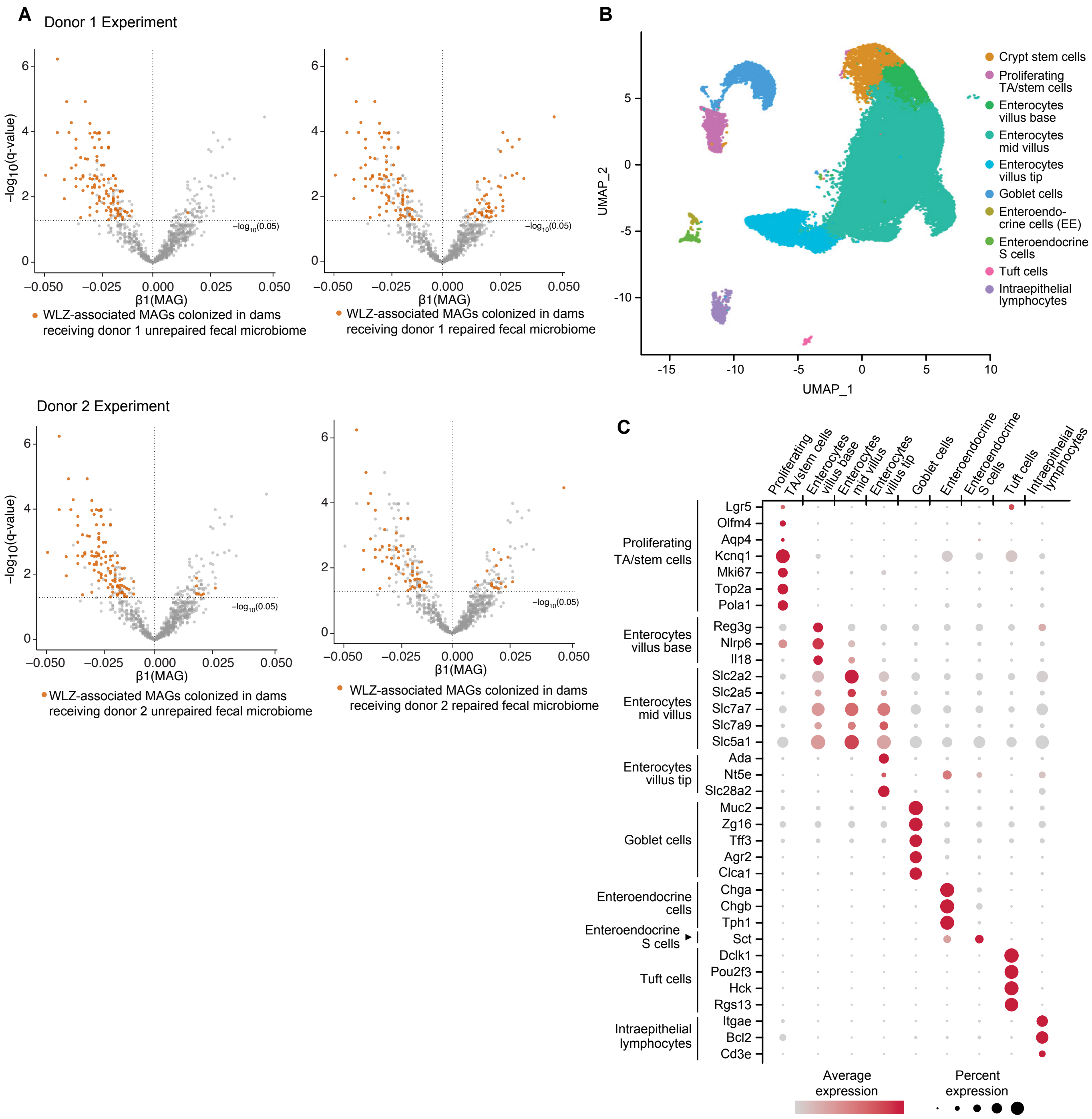

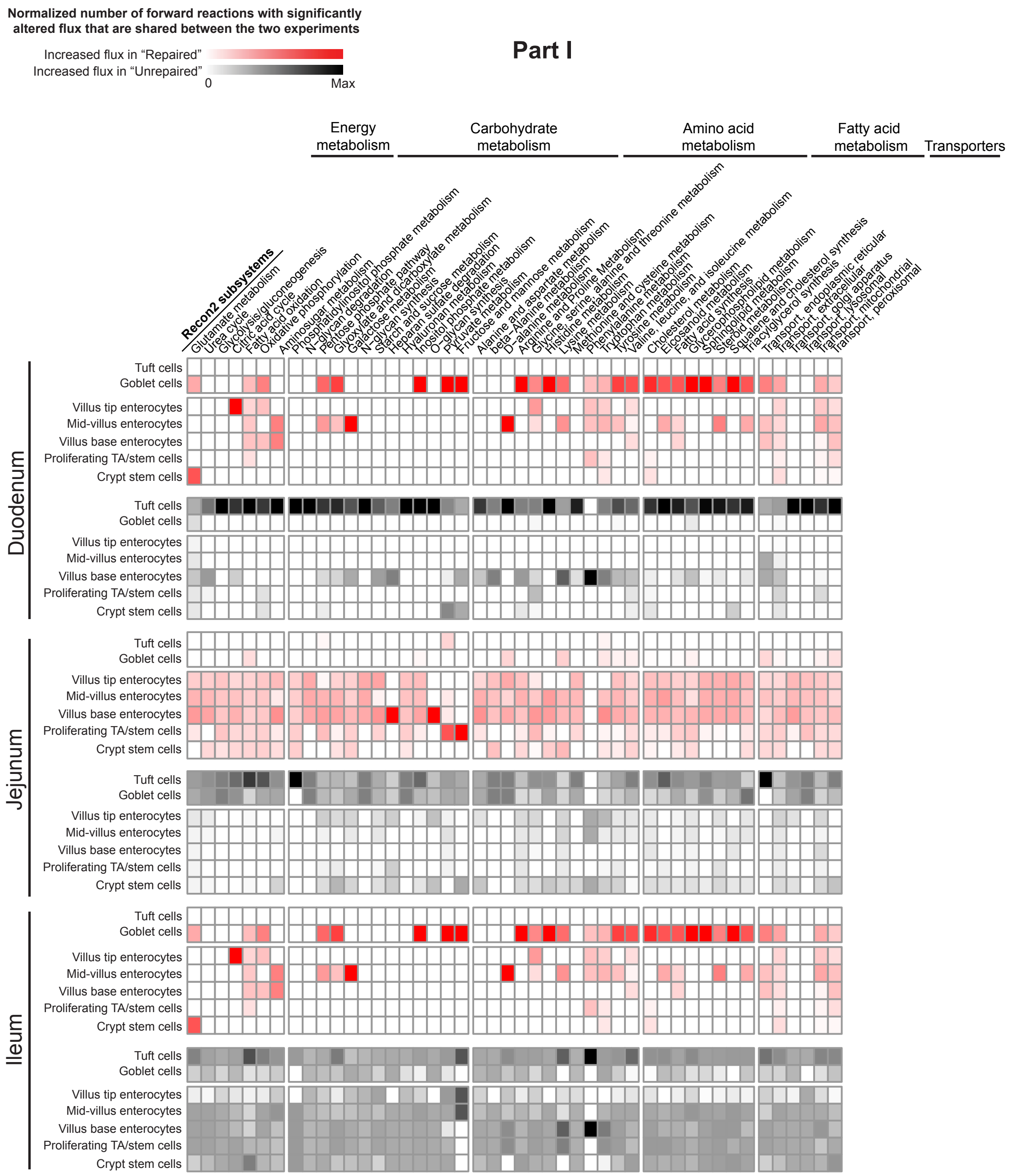

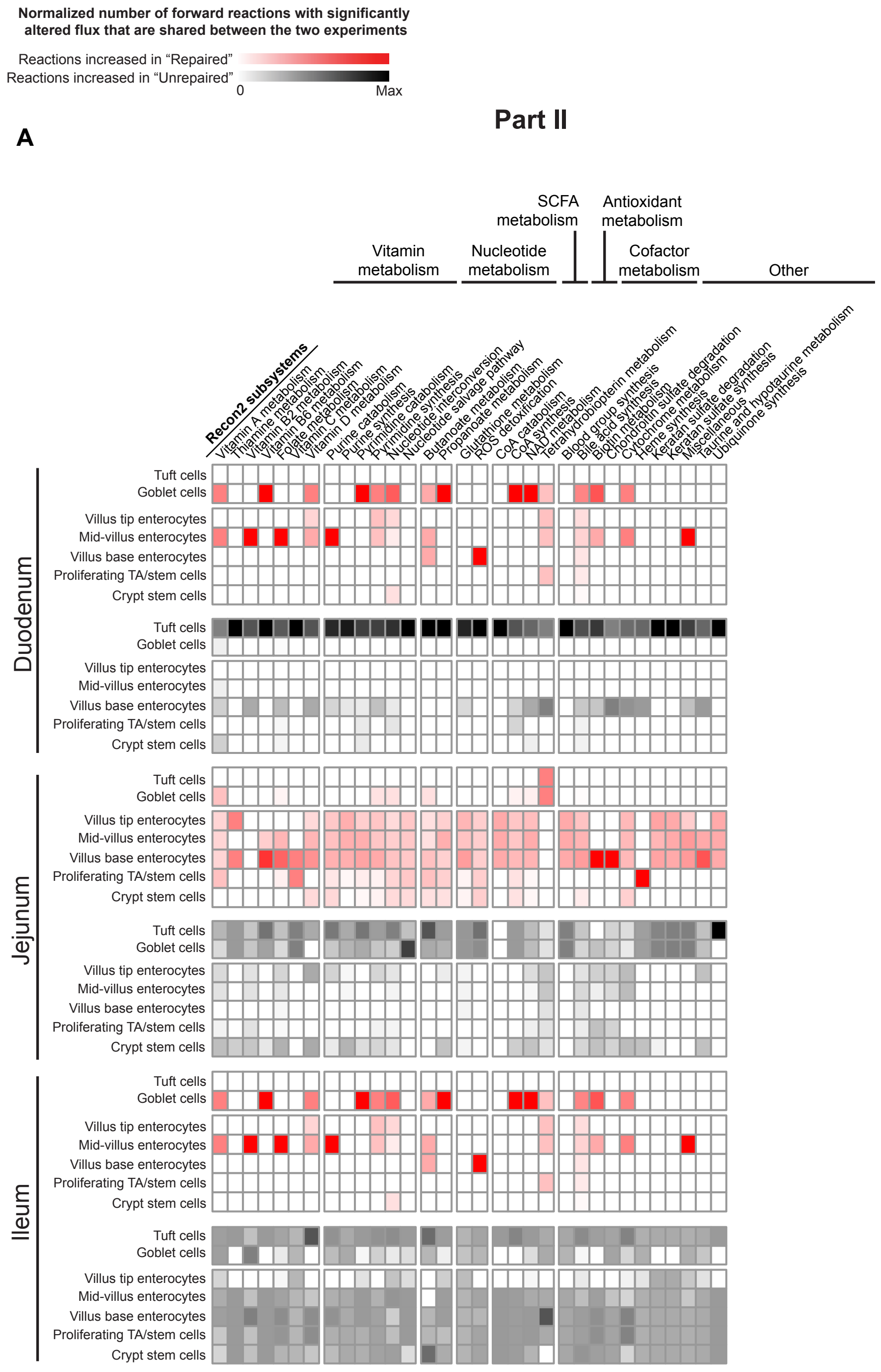

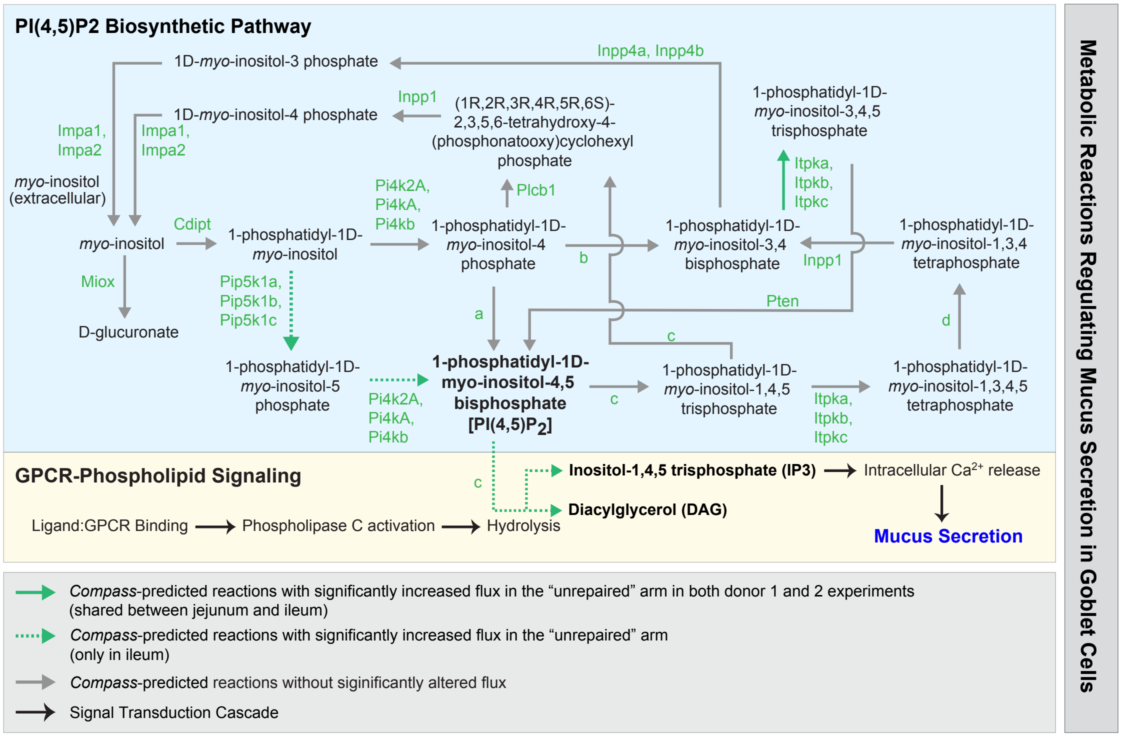

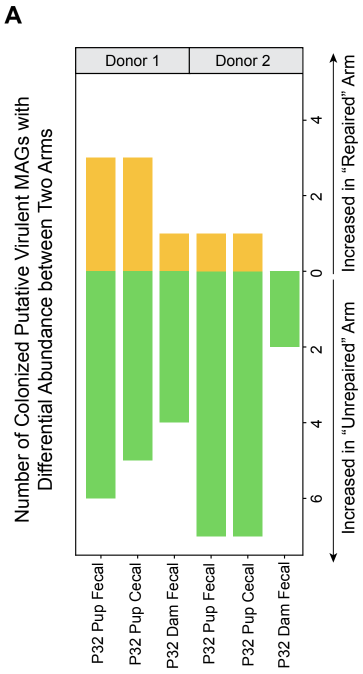

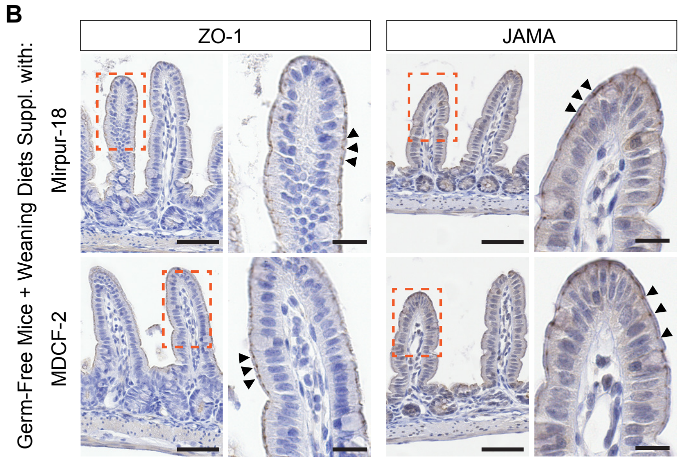
